# Anterior temporal lobes show route-dependent sensitivity to conceptual specificity: Evidence from MEG and EEG

**DOI:** 10.64898/2026.09.28.755013

**Authors:** Ryan M.C. Law, Olaf Hauk, Matthew A. Lambon Ralph

## Abstract

Understanding how the brain represents meaning at different levels of specificity, from general categories (*bird*) through combined phrases (*nocturnal bird*) to specific exemplars (*owl*), is a central question in semantic cognition and its disorders. The anterior temporal lobes (ATL) have been implicated in both activating stored conceptual representations and combining them, yet whether these two routes to conceptual content engage the ATL in the same way remains an open question. We recorded concurrent MEG-EEG from 36 participants reading 100 conceptual triads varying parametrically in specificity: general words (*bird*), adjective-noun phrases (*nocturnal bird*), and specific words (*owl*). Source analyses revealed a non-linear pattern in bilateral ATL (400-520ms): high-specificity concepts elicited greater activity than both low-specificity words and mid-specificity phrases. Bilateral inferior frontal cortex showed this effect with similar timing (430-630ms). N400 amplitude and ATL source activity showed a functional dissociation in their sensitivity to word frequency, ruling out a purely frequency account of the ATL and IFC effects. Results imply that accessing high-specificity concepts requires individuating stored representations among competitors, whereas phrases construct meaning through a different route. These findings extend the unified semantic hub account by revealing that computational operations within the ATL vary depending on task and stimulus demands.

## 1. Introduction

Understanding how the brain constructs meaning at different levels of specificity, from general categories to specific exemplars, is foundational to theories of semantic and language processing and impairments. The neural bases supporting this function remain unclear when meaning is accessed via retrieval through single words or constructed through phrases. While semantic memory (how we represent conceptual knowledge) and semantic composition (how we compose concept knowledge) have been studied separately, both literatures consistently implicate the anterior temporal lobe (ATL). Motivated by computational models proposing a unified semantic hub (Hoffman et al., 2018), we investigate whether the ATL responds similarly when meaning is specified via direct lexical retrieval (*owl*) versus compositional construction (*nocturnal bird*).

The hub-and-spokes (H&S) framework proposes that the ATL serves as a transmodal semantic hub integrating sensorimotor and verbal information from distributed cortical regions into unified conceptual representations (Rogers et al., 2004; Patterson et al., 2007; Lambon Ralph et al., 2017). Evidence for this hub comes from semantic dementia (SD), a syndrome characterised by progressive ATL atrophy/hypometabolism and graded conceptual loss (Warrington, 1975; Schwartz et al., 1979; Snowden et al., 1989; Hodges et al., 1992, 1995; Mummery et al., 1999, 2000; Rogers et al., 2004; Patterson et al., 2006, 2007).

Notably, specific/subordinate concepts (e.g., *canary*) are affected before general/superordinate concepts (e.g., *bird*) (Hodges et al., 1992; Rogers et al., 2015; Rogers & Patterson, 2007; Woollams et al., 2008). Subordinate concepts share highly overlapping features (e.g., has-feathers, can-fly) and have relatively fewer distinctive features (e.g., is-nocturnal, hunts). In contrast, superordinate concepts share a broader set of features that captures similarity relations across subordinate instances. Accordingly as formally demonstrated in the hub-and-spoke computational model, SD patients struggle more with subordinate concepts first because when the hub degrades, it can no longer differentiate subordinate concepts (Rogers et al., 2004, 2015; Rogers & Patterson, 2007).

One prediction of the hub account is that the ATL hub would see increased activity for more specific concepts relative to general concepts, reflecting the greater computational demand of selecting among similar competitors in densely populated regions of conceptual space. This prediction is borne out in functional neuroimaging in healthy adults (Gauthier et al., 1997; Grabowski et al., 2001; Bright et al., 2004; Tyler et al., 2004).

The neural bases of semantic composition have been extensively studied in the recent decades using a minimal two-word phrase reading paradigm (Graves et al., 2010; Price et al., 2015, 2016; Schell et al., 2017; Coutanche et al., 2019; Fritz & Baggio, 2020; Olstad et al., 2020; Graessner et al., 2021; Kochari et al., 2021; Li & Pylkkänen, 2021; Matar et al., 2021; Phillips & Pylkkänen, 2021; Fritz & Baggio, 2022; Murphy et al., 2022; Parrish & Pylkkänen, 2022; Hardy et al., 2023; Zuanazzi et al., 2024; for reviews, see Pylkkänen, 2019b, 2019a; Călinescu et al., 2023). This literature points to the left ATL and, to a lesser extent, the right ATL, for conceptual combination, with increased activity between 200-250 ms post second-word onset for combinatory items (e.g., *red boat*), relative to non-combinatory controls (e.g., *cup, boat* or *xtq, boat*) across language production and comprehension (Pylkkänen, 2019a, 2019b). Crucially, Westerlund and Pylkkänen (2014) showed that ATL composition effects are modulated by noun specificity: adjective-noun phrases elicited increased ATL activity only when the noun was general (*blue boat*), not when it was already specific (*blue canoe*), suggesting the ATL is sensitive to both compositional operations and the specificity of the resulting concept.

The functional-anatomical overlap between semantic memory and semantic composition in the ATL implies a shared substrate for both systems (“the unified hub hypothesis”). This idea finds computational and empirical support. Hoffman et al. (2018) showed that a single hidden hub layer trained on both linguistic context and sensorimotor features can reproduce both SD deficits and context-sensitive linguistic interpretations, providing a computational framework for the unified hub hypothesis. Empirical support comes from Law et al. (2026), who crossed semantic memory variables (noun concreteness: *lettuce* vs. *fiction*) with compositional variables (adjective type: *bad* vs. *fake*) and found bilateral ATL sensitivity to both, suggesting a common representational substrate. However, that study did not address whether the ATL responds similarly when conceptual content is approximately matched but the route to that content differs.

The current study addressed this gap by creating 100 conceptual triads, each instantiating three levels of specificity: a general single word (*bird*), a compositionally specified phrase (*nocturnal bird*), and a specific single word (*owl*). The semantic memory and composition literatures show that the ATL is sensitive to conceptual specificity, but the form of this relationship remains unclear when specificity is varied both within and across words. Two possibilities emerge from the literature:

Possibility 1 (graded specificity): If the ATL tracks conceptual content in a graded manner regardless of access route, neural activity should scale monotonically with specificity: Low < Mid < High, whether concepts are activated through single words or composed through phrases.

Possibility 2 (lexical individuation): If activating specific concepts imposes distinct computational demands (perhaps because subordinate concepts must be differentiated from many similar competitors), we might observe elevated ATL activity for high-specificity lexical items (*owl*) but not for compositionally derived intermediate specificity (*nocturnal bird*), even if the latter approximates the former in semantic content.

We recorded concurrent MEG and EEG data from 36 participants in a word and phrase comprehension task and analysed source-localised neural activity in bilateral ATL and semantic control regions using mass-univariate cluster-based permutation tests.

## 2. Methods

The study was approved by the local ethics committee. The study was pre-registered on the Open Science Framework on July 14, 2023, before data collection began (link withheld for review). Data from the same sessions were also used in a separate, previously published study (Anonymous, 2026).

### 2.1 Participants

A cohort of 36 adults (age range 18-40 years, M = 26 years, SD = 6 years; 27 women) participated in the MEG-EEG study. To ensure sufficient statistical power to detect a medium-sized effect (effect size *f* = 0.25, *α* = 0.01, *β* = 0.05), the sample size was determined *a priori* using *G*Power* (v3.1.9.6; Faul et al., 2007). All recruited individuals were right-handed, native English speakers with normal or corrected-to-normal vision and reported no history of neurological disorders. Participants provided informed consent in accordance with the local ethics committee and received financial compensation for their time.

### 2.2 Stimuli

The stimulus set was constructed around 100 conceptual ‘triads’ (total n=300), each designed to instantiate three distinct levels of specificity: low, mid, and high.

The nouns (*e.g., bird, owl*) denoted concrete concepts. These were sampled from the Brysbaert et al. (2014) norms, restricted to items with high concreteness ratings (4.0-5.0), known by >90% of raters, and consisting of 3-10 letters. The mid-specificity condition (e.g., *nocturnal bird*) consisted of two-word phrases created by selecting an adjective that adds a key conceptual feature to the low-specificity noun, thereby making its meaning more similar to the high-specificity noun. For the single-word baseline trials (low and high), the first stimulus was a length-matched, unpronounceable consonant string (*e.g., qtpv*) to serve as a non-semantic visual placeholder.

We adopted a parametric design with three levels of specificity rather than a fully crossed 2x2 factorial design (e.g., crossing Specificity with Composition). While a full factorial design would allow for a direct comparison of composition across specificity levels, the combination of a high-specificity noun with a restrictive modifier (e.g., *nocturnal owl*) often results in redundancy or unnatural phrasing, as the modifier’s features are typically already inherent to the specific concept. Our primary objective was to chart the correlates of a clear, graded conceptual progression from a general category (*bird*), through a specified description (*nocturnal bird*), to a specific lexical entry (*owl*), a trajectory that has not yet been empirically studied in the literature. Consequently, we prioritised this goal over a balanced factorial structure, accepting that the Mid condition differs from the Low and High conditions in its compositional structure (adjective-noun phrase vs. single word) to ensure the validity of the specificity gradient.

In prioritising the conceptual relations within each triad, we acknowledge that a perfect matching of psycholinguistic variables (e.g., frequency, age of acquisition) relevant for conceptual processing was not always feasible (see Table 1, Zipf frequency difference between Low and High conditions). We took this into account below when we interpreted the pattern of our results. Table 1 summarises the distribution of a range of psycholinguistic variables of our target nouns across conditions.

**Table 1.** Psycholinguistic characteristics of the nouns used in the stimulus set. Low and Mid items share the same nouns.

|  | Low | Mid | High |
| --- | --- | --- | --- |
| **Concreteness rating** | 4.65 (0.41) | 4.65 (0.41) | 4.71 (0.35) |
| **Imageability** | 6.32 (0.62) | 6.32 (0.62) | 6.45 (0.50) |
| **Length (number of letters)** | 5.69 (1.88) | 5.69 (1.88) | 6.05 (1.85) |
| **Zipf frequency** | 4.44 (0.66) | 4.44 (0.66) | 3.85 (0.65) |
| **Age of acquisition (years)** | 5.59 (2.10) | 5.59 (2.10) | 6.5 (2.14) |
| **Valence** | 5.66 (0.95) | 5.66 (0.95) | 5.53 (0.93) |
| **Lexical decision latency (ms)** | 548.38 (39.43) | 548.38 (39.43) | 572.38 (50.17) |

Mean values for each measure are reported, with SD in parentheses.

**Table 2.**
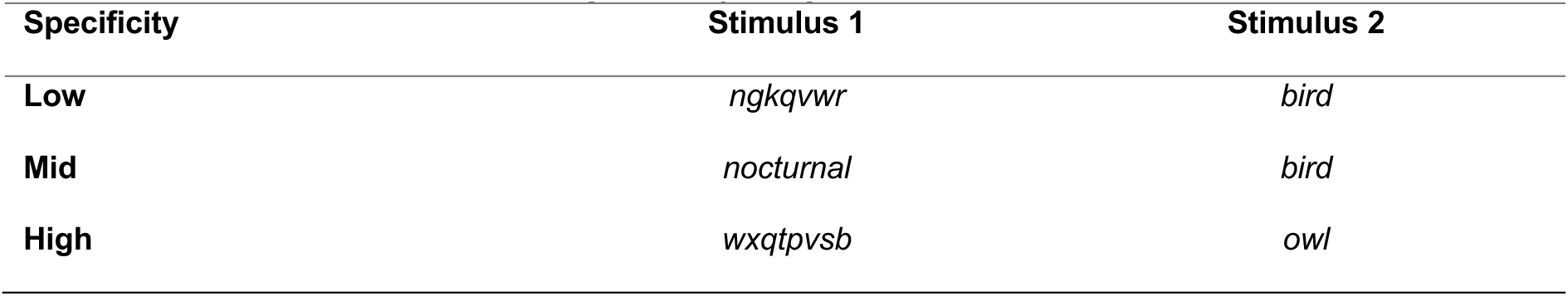
An illustrative set of items showing the study design.

### 2.3 Stimulus Validation

#### 2.3.1 Behavioural Norming

To ensure that our stimulus triads followed the intended parametric increase in specificity, we conducted a behavioural norming study with a separate sample of participants. A total of 52 participants (recruited via Prolific (www.prolific.com) and hosted on the Gorilla (www.gorilla.sc) completed a forced-choice specificity rating task. On each trial, participants were presented with two items from a triad (e.g., bird and nocturnal bird) and were asked to select the item that was more specific or general (blocked and counterbalanced). Participants were given opportunity to practice with feedback in blocks of 9 trials until they were ready to proceed. Each participant completed 300 trials.

Analysis of sensitivity (d’) using a linear mixed-effects model (subject as random effects) revealed significant differences between specificity conditions. Sensitivity (d’) was significantly lower for the *High-Mid* condition (M = 1.72), compared to both *Low-Mid* (M = 2.65), M diff = −0.93, 95% CI [−1.21, −0.64], *p* < .001, and *High-Low* (M = 2.66), M diff = −0.94, 95% CI [−1.22, −0.65], *p* < .001. The latter two did not differ, M diff = 0.010, 95% CI [−0.27, 0.29], *p* > .9. These behavioural results hinted at the possibility that direct lexical access to specific concepts (e.g., *owl*) involves distinct cognitive demands compared to phrasal description (e.g., *nocturnal bird*). See Supplementary Figure 1A.

Our specificity manipulation also engendered significant differences in reaction times (RT), as revealed by a linear mixed-effects model (subject as random effects). Post-hoc Tukey comparisons indicated that distinguishing a phrase from a specific noun (*Mid-High*) was significantly slower (M = 1.88 s) than distinguishing a general noun from a phrase (*Low*-Mid) (M = 1.48 s), M diff = 0.41 s, 95% CI [0.19, 0.62], *p* < .001, and significantly slower than distinguishing a general noun from a specific noun (*High-Low*) (M = 1.63 s), M diff = 0.21 s, 95% CI [0.04, 0.46], *p* < .001. The latter two did not differ, M diff = 0.16, 95% CI [−0.06, 0.37], *p* > .1. See Supplementary Figure 1B.

#### 2.3.2 Computational Validation

We also sought to quantify specificity using two computational metrics. First, to verify taxonomic depth, we calculated the maximum hypernym depth in WordNet (Miller, 1994) for our single-word items (i.e., low and high specificity items). Our stimulus set contained 96 unique low-high specificity pairs. A paired t-test revealed that high-specificity words were significantly deeper in the semantic hierarchy than low-specificity words, M = 1.58 levels, *t*(95) = 6.58*, p* < .001.

Second, to characterise the distributional relationship between conditions, we extracted vector embeddings using Sentence-BERT (Reimers & Gurevych, 2019). We calculated, for each conceptual triad, a specificity gain score: the cosine similarity between the phrasal condition (*nocturnal bird*) and the high-specificity target (*owl*) minus the cosine similarity between the low-specificity baseline and the same high-specificity target (e.g., *bird* vs. *owl*). A value of 0 would indicate that the phrase (*nocturnal bird*) is as close to the specific target (*owl*) as the general word (*bird*) is in the sBERT vector space; positive values indicate the phrase has moved toward the high-specificity target.

A one-sample *t*-test revealed a small but significant decrease in cosine similarity due to composition (i.e., *nocturnal bird* < *owl*), M = −0.024, *t*(99) = −2.16, *p* < .033. This indicates that composition (*nocturnal bird*) produced a representation that is distributionally more distant from the high-specificity target (*owl*) than the bare noun (*bird*) alone. This implies that composed representations are not “mere” approximations of high-specificity conceptual representations.

### 2.4 Experiment Procedure

Participants were instructed to read the stimuli silently for comprehension. The 300 trials were fully randomised and presented across five experimental blocks.

The sequence of events in each trial unfolded as follows (Figure 1). A central fixation cross (300 ms) was replaced by the first stimulus (an adjective or a consonant string), which was displayed for 300 ms. After a 300 ms blank interval, the second stimulus (the noun) appeared for 300 ms, followed by a final 500 ms blank period. To ensure sustained attention, a ‘yes/no’ comprehension question appeared on 10% of trials, requiring a button-press response. Each trial concluded with an inter-trial interval that was jittered around an average of 1.5 s.

**Figure 1.**
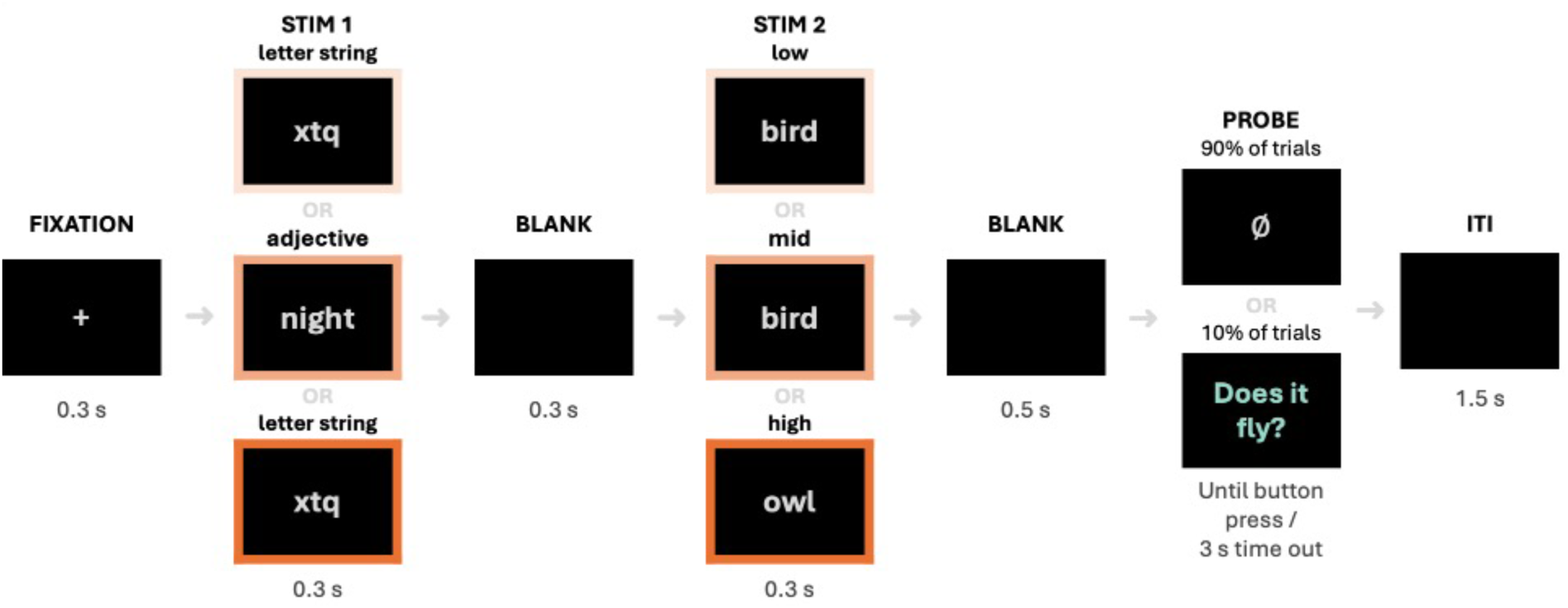
Trial structure. Task probes were included for 10% of trials to encourage comprehension and attention.

**Figure 2.**
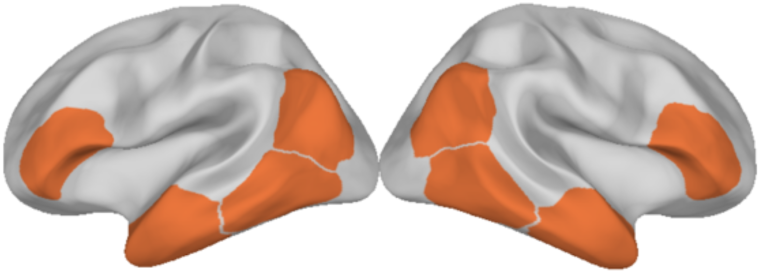
Regions of interest.

### 2.5 Data Acquisition and Preprocessing

Concurrent MEG and EEG recordings were acquired in a magnetically shielded room using a 306-channel TRIUX neo system (MEGIN, Espoo, Finland). The MEG system comprised 204 planar gradiometers and 102 magnetometers. EEG was recorded from a 64-electrode EEG cap (EasyCap GmbH, Herrsching, Germany) in a standard 10-20 layout, with a nose reference. Vertical and horizontal electrooculograms (EOG) were monitored for eye movements. Impedance levels were reduced to 10 kOhms or lower during preparation when possible. Prior to recording, each participant’s head shape and the locations of HPI coils and fiducial landmarks were digitised for accurate co-registration with their individual anatomical MRI scans. All data were sampled at 1 kHz with an online high-pass filter of 0.03 Hz. As for MR images, for each participant, a T1-weighted structural scan was acquired using a Siemens 3T Prisma scanner.

The raw data were processed using the MNE-Python (v1.4.2; Gramfort et al., 2013, 2014). An offline 0.1-40 Hz bandpass filter was first applied, using a finite impulse response filter design implemented with a Hamming window method. For MEG data, bad and flat channels were automatically detected using “find_bad_channels_maxwell” and supplemented by visual inspection. Continuous head position was estimated from the HPI coil signals. We then applied spatiotemporal Signal Space Separation (stSSS) (Taulu & Kajola, 2005) using “maxwell_filter”, incorporating fine calibration and cross-talk correction. To compensate for head movement, all data were transformed to a common reference head position taken from the block in the middle of the experiment (i.e., block 3). For EEG data, noisy channels were interpolated using the spherical spline method (Perrin et al., 1989), and ocular artefacts were identified via Independent Component Analysis (ICA) correlated with EOG channels and subsequently removed.

The pre-processed continuous data were then segmented into epochs spanning from 200 ms before the fixation cross to 1.4 s after the first word onset. A common pre-fixation baseline (−200 ms to 0 ms relative to fixation) was used for baseline correction before the data were down-sampled to 250 Hz and subjected to an automated artefact rejection procedure (“autoreject”, Jas et al., 2017). A dataset from one participant were particularly noisy and were excluded from subsequent analyses. Around 80 trials per condition entered further analyses.

### 2.6 Univariate ERP Analysis

To analyse ERP responses to the noun, we conducted analyses in two separate time windows, which have been previously associated with semantic composition: N400 (Neufeld et al., 2016) and P600 (Fritz & Baggio, 2020). Difference waves were computed for each participant by subtracting the low-specificity evoked response from the high-specificity and mid-specificity evoked responses respectively, and the mid-specificity response from the high-specificity response. The difference waves were then averaged across a centroparietal electrode cluster (C1, Cz, C2, CP1, CPz, CP2, P1, Pz, P2) and across the two time windows: 350–500 ms and 600-800 ms post noun onset. This yielded one measurement per time window per condition per participant. Prior to testing, normality was assessed using the Kolmogorov-Smirnov test. Where normality held, one-sample t-tests against zero were used; where it was violated, Wilcoxon signed-rank tests were used instead. All tests were one-tailed, testing for a negative deflection (more negative than zero).

### 2.7 Source Reconstruction

We used *FreeSurfer* (v6.0.0; Fischl, 2012) alongside the Python package *MNE-Python* (v1.4.2; Gramfort et al., 2013, 2014) to reconstruct subject-specific anatomical models of the scalp, inner/outer skull, and cortical surface. T1-weighted anatomical MR images were first segmented using the automated segmentation algorithms in *FreeSurfer* (Dale et al., 1999) to obtain the reconstructed scalp surface.

Forward and inverse solutions were calculated using *MNE-Python*. We used a three-layer boundary element (BEM) model to compute the forward model including each participant’s scalp, outer skull surface, and inner skull surface (conductivity parameters are 0.3, 0.006, and 0.3 respectively). The MEG-EEG sensor configurations and MR structural images were then co-registered by matching the scalp digitisation points to the scalp surface. Subject-specific source spaces contained 4098 dipole locations in each hemisphere. To calculate the inverse solution, we estimated the noise covariance matrix using the 200 ms window prior to fixation and then regularised it using an automated model selection method (Engemann & Gramfort, 2015). The regularised noise covariance matrices and forward solutions were used to assemble the inverse operator using L2 minimum norm estimation (L2-MNE) (Hauk et al., 2022) with a loose orientation constraint value of 0.2 and depth weighting of 0.8. The inverse operator was then regularised assuming a signal-to-noise ratio of 3. After applying the inverse operator to sensor evoked responses, individual-specific cortex-wide source estimates were morphed to a common space (‘*fsaverage’*) for group analyses.

### 2.8 Mass-univariate region-of-interest (ROI) analyses

While our primary hypotheses concern the role of the ATL in unifying semantic memory and composition, semantic cognition is supported by a network of brain regions (Lambon Ralph et al., 2017). To test the anatomical specificity of our predicted effects and to situate the ATL’s function within the broader neurocognitive architecture of semantics, we also included a set of key control regions. We defined ROIs for the temporo-parietal cortex (TPC), another critical node implicated in conceptual processing during phrase and sentence comprehension (Matchin et al., 2019; Pallier et al., 2011), to assess the generality of our findings across the core semantic network. Furthermore, we included the inferior frontal cortex (IFC) and posterior temporal lobe (PTL), regions robustly associated with the semantic control network (Jackson et al., 2021; Lambon Ralph et al., 2017). The inclusion of these control regions allows us to dissociate neural activity related to the representation of conceptual meaning (hypothesised to be centred on the ATL) from semantic control.

Our ROIs were defined anatomically based on prior literature. First, we identified four MNI coordinates (Table 3) from foundational studies and meta-analyses of semantic cognition (Lambon Ralph et al., 2017; Jackson, 2021) and phrase comprehension (Bemis & Pylkkänen, 2011; Pallier et al., 2011; Pylkkänen, 2019a, 2019b). Second, around each of these seed coordinates, we grew a spherical label with a 30-mm radius (40-mm for the ATLs). Finally, to ensure bilateral coverage, we included the corresponding right-hemisphere homologues by using the negative x-coordinate of each seed point.

**Table 3.** MNI coordinates of ROIs.

| ROIs | MNI coordinates | ROIs | MNI coordinates |
| --- | --- | --- | --- |
| Left ATL | [-57, -2, -33] | Right ATL | [57, -2, -33] |
| Left PTL | [-50, -52, -6] | Right PTL | [50, -52, -6] |
| Left IFC | [-45, -66, 24] | Right IFC | [45, -66, 24] |
| Left TPC | [-51, 30, 6] | Right TPC | [51, 30, 6] |

To analyse the time course of neural activity in each ROI, we employed non-parametric cluster-based permutation tests (Maris & Oostenveld, 2007), implemented in the Python package *Eelbrain* (version 0.39; Brodbeck et al., 2023). The analysis pipeline began by averaging cortical activity intensity across all sources within an ROI to produce a single time-series. We then fitted statistical models to this time course across the entire 800 ms post-stimulus window, with subjects treated as a random effect. To investigate our specific research questions, we used one-way ANOVAs with a fixed effect for specificity of three levels (low, mid, high). We quantified the effect size of any significant temporal clusters using Cohen’s *f*. The statistical significance of these clusters was estimated via 5,000 Monte Carlo simulations. Finally, to account for multiple comparisons across the eight ROIs, we applied a False Discovery Rate (FDR) correction using the Benjamini-Hochberg procedure (Benjamini & Hochberg, 1995) and reported the resulting corrected *q*-values.

### 2.9 Behavioural Analyses

While comprehension probes were included to ensure attentional engagement, they were not controlled for length or difficulty. Accordingly, the following behavioural analyses are reported for completeness and are not central to our primary conclusions.

Statistical analyses were performed using the lme4 (Bates et al., 2015) and lmerTest (Kuznetsova et al., 2017) packages in R. Two participants and one item were excluded from analysis due to poor performance. Reaction times (RT) for correct trials were analysed using a linear mixed-effects model. Prior to analysis, outliers were trimmed by removing RTs exceeding 2.5 standard deviations from each participant’s mean. Accuracy was analysed using a generalised mixed-effects logistic regression. Both models included specificity (low, mid, high) as a fixed effect. Random effects structures were initially maximal, including by-participant random slopes for specificity. To achieve convergence, structures were simplified to include only by-participant and by-item random intercepts for the modelling of accuracy; As for reaction times, structures included by-participant random slopes for specificity. Model fit was assessed using marginal and conditional R^2^ (MuMIn package).

## 3. Results

### 3.1 Behavioural Results

As aforementioned, comprehension probes engaged comprehension and attention but were not matched for psycholinguistic variables. Accordingly, behavioural data are only reported descriptively. Overall, accuracy was high across all conditions, low = 94%, mid = 93%, high = 95%, and reaction times (seconds) were comparable across conditions, low = 1.36, mid = 1.41, high = 1.34, confirming that participants remained attentive throughout the task.

We found no main effects of specificity on accuracy, *χ*^2^(2) = 0.81, *p* = .67, or reaction times, *χ*^2^(2) = 1.12, *p* = .57. Model fit metrics disclosed that while the fixed effect of specificity explained negligible variance (Marginal *R^2^*: RT = .005, Accuracy < .008), the overall models accounted for a substantial portion of the variance (Conditional *R^2^*: RT = .57, Accuracy = .22). These results indicate that task demand or difficulty are unlikely confounds for our neural findings.

### 3.2 ERP Effects of Specificity

After the presentation of the first stimulus, we see a larger N400 amplitude for mid specificity condition (e.g., *nocturnal*) relative to low and high specificity conditions (e.g., *ngkqvwr*) (Figure 3). The presence of N400 effects, consistent with the N400 literature (Kutas & Federmeier, 2011), confirmed that participants were engaged during the experiment and were processing stimuli at the conceptual level in a simple comprehension task.

**Figure 3.**
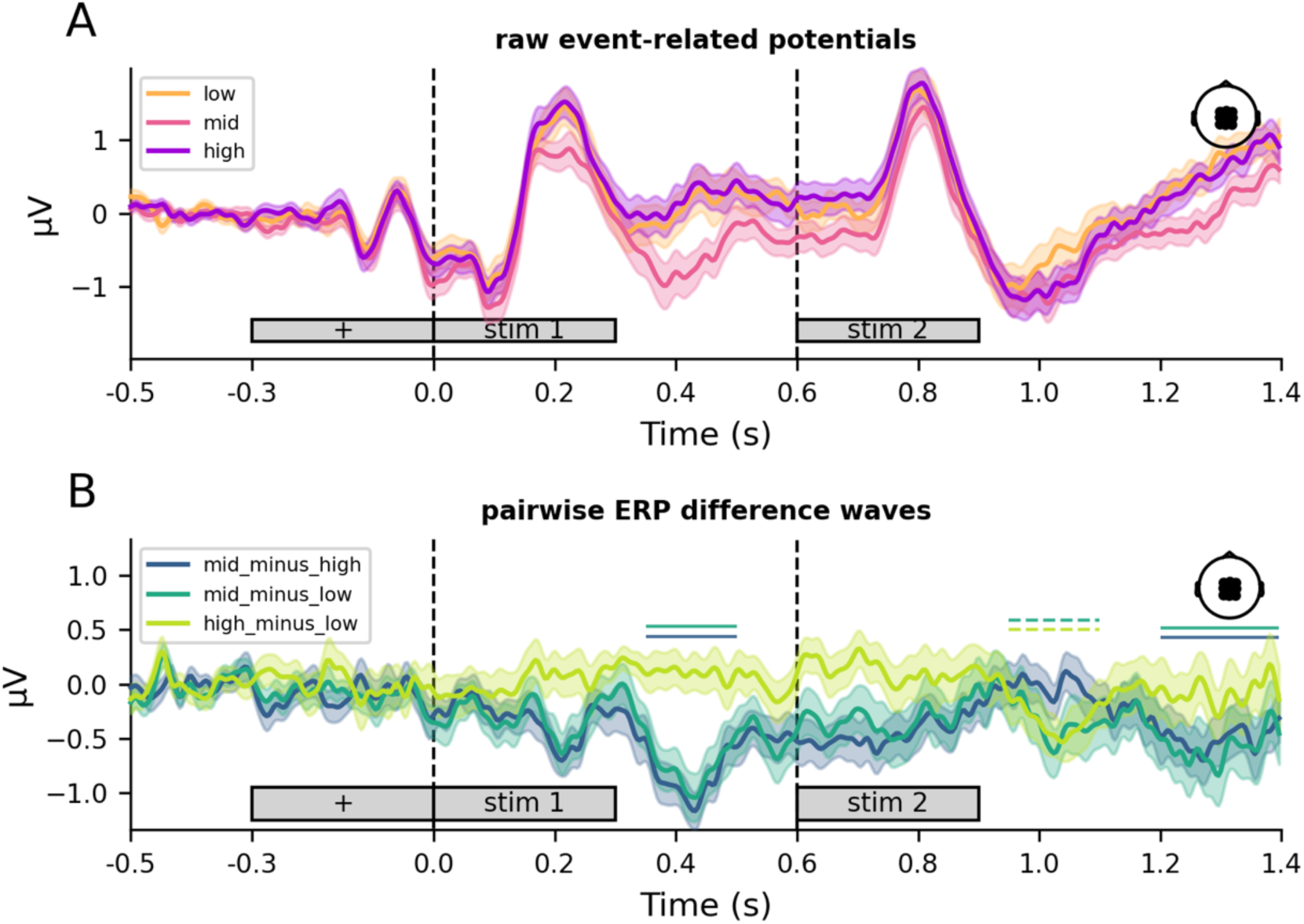
Event-related potentials of different specificity levels. (A) Raw grand-average ERPs time-locked to the onset of the first stimulus at centroparietal electrodes. Coloured inset legend (right) plots the locations on the EEG cap. Shaded region represents one standard error of the mean. (B) ERP difference waves each representing one contrast. Horizontal bars above each waveform indicate t-test results. Bar colour corresponds to the contrast as indicated in the legend. Solid bars indicate effects that survive FDR correction (p_corrected < .05); dashed bars indicate trends that did not survive FDR correction.

Our analyses focused on neural responses elicited by the nouns. Group-level one sample *t*-tests revealed that, relative to low-specificity controls, neural responses were more negative in centroparietal electrodes at the N400 time window (350-500 ms post noun onset) for both high, *t*(34) = −1.76, *p* = .044, and mid specificity items, *t*(34) = −1.60, *p* = .060 (Figure 3) , though neither survived FDR correction (*p* = .078 and *p* = .089 respectively). At a later time window, between 600-800 ms post noun onset, neural activity from the same electrodes was more negative for mid specificity items (i.e., phrases) relative to both low, *t*(34) = −2.17, *p* = .019, and high, *t*(34) = −2.48, *p* = .009, specificity items (i.e., single words), both of which survived FDR correction (*p* = .042 and *p* = .028 respectively).

### 3.3 Main Effects of specificity in the anterior temporal lobes and inferior frontal cortices

Cluster-based permutation tests revealed a main effect of specificity, *p =* .010, in bilateral ATLs (Figure 4). Clusters corresponding to this effect were localised at around 400-520 ms post-noun-onset, with high specificity items eliciting higher activity relative to low and mid specificity items (Cohen’s *f* = 0.15).

**Figure 4.**
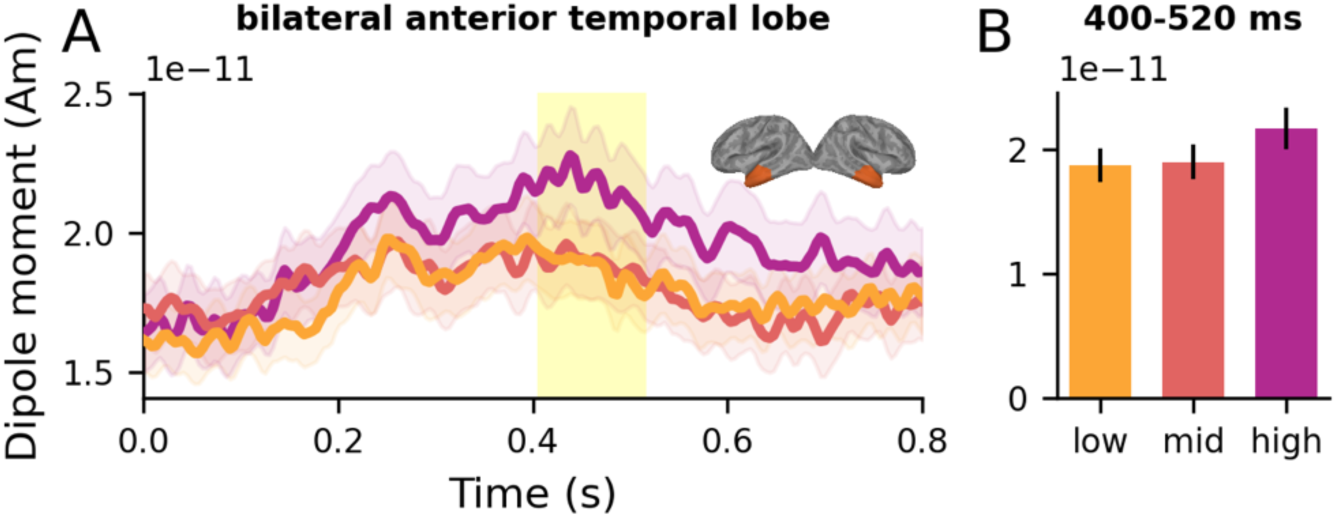
Main effect of specificity across left and right anterior temporal lobes. Time courses time locked to onset of the second stimuli represent group averages of each condition and error represents one within-subjects standard error of the mean. Yellow vertical spans represent temporal clusters identified in cluster-based permutation tests.

To test the spatial specificity of our predicted effects, we also tested other ROIs (see Methods). Our analyses revealed main effects of specificity (*p* = .002) and hemisphere (*p* = .009) across left and right IFC. There were no significant interactions between hemisphere and specificity. Specificity effects were associated with a temporal cluster around 430-630 ms post noun onset (Cohen’s *f* = 0.16; Figure 5A,B), driven by elevated activity for high specificity items. As for the effect of hemisphere, a temporal cluster was identified at around 270-440 ms post noun onset (Cohen’s *f* = 0.16; Figure 5C,D).

**Figure 5.**
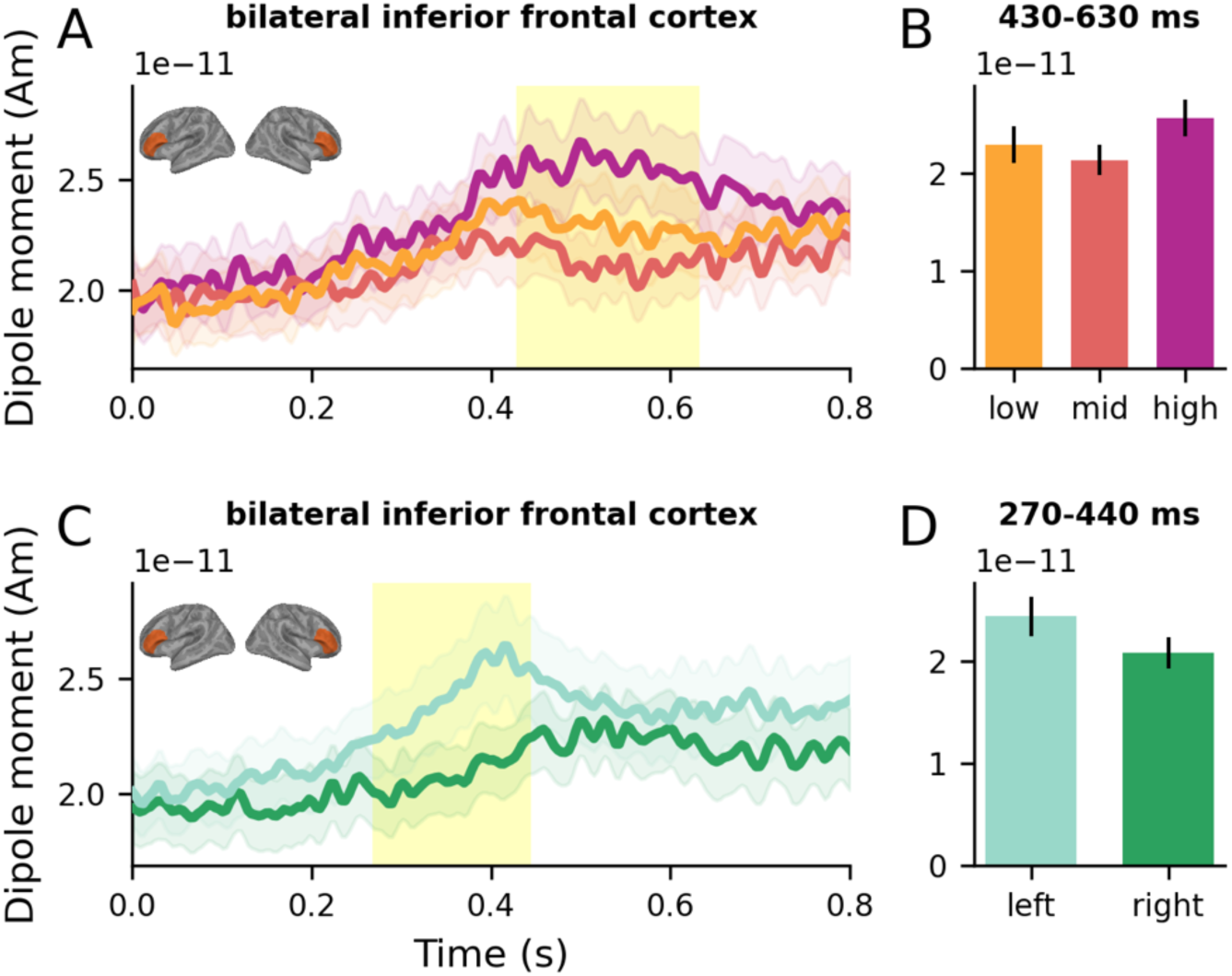
Main effects of specificity (A,B) and of hemisphere (C,D) across left and right inferior frontal gyri. Time courses represent group averages of each condition and error represents one within-subjects standard error of the mean. Yellow vertical spans represent temporal clusters identified in cluster-based permutation tests.

Other tested ROIs did not show significant effects of specificity. For individual ROI time series, see Supplementary Figure 2. For uncorrected *t*-maps of activity between high and low specificity words, see Supplementary Figure 3.

## 4. Discussion

Understanding how the brain represents meaning at different levels of specificity, from general categories to specific exemplars, is foundational to theories of semantic cognition. The hub-and-spokes framework proposes that the anterior temporal lobes (ATL) serve as a transmodal semantic hub for conceptual representation, yet the neural bases supporting conceptual specification remain unclear when meaning is accessed via different routes: direct lexical retrieval versus compositional construction. We addressed this question using a parametric design in which conceptual specificity varied across both single words (*bird*, *owl*) and phrases (*nocturnal bird*), allowing us to test whether ATL activity tracks conceptual content regardless of access route (graded specificity; Possibility 1) or whether lexical and compositional routes engage the hub differently (lexical individuation; Possibility 2).

Our findings support the lexical individuation account. Rather than a monotonic increase with specificity (low < mid < high), bilateral ATL showed a non-linear, step-like pattern: high-specificity single words (*owl*) elicited reliably greater activity (400-520ms post-noun-onset) than *both* low-specificity words (*bird*) and mid-specificity phrases (*nocturnal bird*), with the latter two conditions statistically indistinguishable. This pattern suggests that accessing fine-grained concepts places enhanced computational demands on the semantic hub compared to compositional phrases.

This pattern is consistent with the notion that activating fine-grained concepts places considerable demands on the semantic hub. According to the H&S framework, the hub must represent concepts in a way that preserves similarity structure and distinctiveness amongst related concepts (Rogers et al., 2004, 2015). High-specificity concepts like *owl* require individuated representations to distinguish them from highly similar competitors (e.g., other birds like canaries, parrots and so on). This demand for fine-grained differentiation among closely related concepts has been consistently observed in semantic dementia, where subordinate-level concepts are disproportionately impaired relative to general categories (Hodges et al., 1992; Rogers et al., 2015; Rogers & Patterson, 2007; Woollams et al., 2008), and in neuroimaging studies showing elevated ATL activity for specific relative to general concepts (Gauthier et al., 1997; Grabowski et al., 2001; Bright et al., 2004; Tyler et al., 2004). The sustained and elevated ATL response for high-specificity items may reflect the computational cost of activating and individuating these representations – as shown previously in computational semantic models (Rogers & Patterson, 2007).

Crucially, compositionally deriving an intermediate level of specificity (*nocturnal bird*) appears to engage the hub differently than directly activating a subordinate concept (*owl*). One possibility is that in adjective-noun phrases, the adjective provides context that constrains the meaning of the upcoming noun. This compositional process creates a more specific conceptual representation than the head noun alone (e.g., *nocturnal bird* is more specific than *bird*), but it may not require the same processes engaged when accessing a highly specified lexicalised entry, or at least not to the same extent because the hub representation has already been “pushed” towards the correct, more specific interpretation by the adjective (somewhat like the effect of semantic cues simulated in the unified computation model of Hoffman and colleagues; Hoffman et al., 2018).

This result provides a new benchmark for computational models of the semantic hub. Hoffman et al. (2018) proposed a framework in which a single set of hub units learn from both sensorimotor features and linguistic context. Our results suggest that future computational work should account for why direct lexical access to high-specificity items may be more costly than composition, producing the step-like increase we observed rather than a continuous gradient.

Our results point to a broader, more bilateral network than the left-lateralised effects reported previously. The engagement of both left and right ATLs is consistent with proposals that ascribe functional collaboration between the ATLs (Patterson et al., 2007; Schapiro et al., 2013; Lambon Ralph, 2014; Jung & Lambon Ralph, 2016). The sustained, bilateral IFC activity further suggests the recruitment of the semantic control network (Lambon Ralph et al., 2017), which is engaged to select and maintain appropriate representations.

Interestingly, these regions showed different peak latencies. ATL effects were observed around 400-520 ms post noun onset, while IFC effects were slightly later at around 430-630 ms. This latency difference may hint at information flow between regions. Future work could explore the precise nature of the interaction between representation and control systems through, for example, effective connectivity analyses. Nonetheless, the coactivation of ATLs and IFCs in our study underscores their joint contribution during conceptual processing. Further, it suggests that task changes can reveal how the core semantic representation hub (ATLs) may dynamically interact with control regions (IFCs) across both hemispheres to support conceptual understanding, opening new questions about the role of semantic control in even basic forms of phrasal meaning construction.

A potential concern is that the elevated ATL response for high-specificity items (*owl*) might reflect psycholinguistic confounds such as word length, frequency, or age of acquisition, rather than conceptual specificity *per se*. However, several features of our design and results argue against these alternative interpretations.

First, if the ATL tracked conceptual rarity, the mid-specificity condition should have elicited the strongest response, as phrasal expressions are not stored as lexical entries and therefore lack the kind of direct frequency-based representation that single words have. The absence of such an effect argues against a frequency-of-concept account.

Second, while the high-specificity condition consisted of words with lower word frequency than the low-specificity condition, the dissociation between our N400 and ATL results argues against a pure word-frequency account. The N400 component, a classic marker of frequency and expectancies, differentiated between the low- and mid-specificity conditions (Figure 3). In contrast, the ATL responses were statistically indistinguishable (Figure 4). This functional dissociation suggests that the ATL effect is specific to the demands of conceptualising a highly specific item, a process that is qualitatively distinct from the continuous variables of frequency or length.

An important finding of our study was the absence of the early left ATL composition effect. In our key comparisons, the mid-specificity phrasal condition (e.g., *nocturnal bird*) elicited a neural response in the ATLs that was statistically indistinguishable from the low-specificity single-word condition (e.g., *bird*). The only factor that robustly modulated ATL activity was high lexical specificity, with single words like *owl* producing a greater response than both the general word and the phrase. This pattern stands in contrast with a large body of prior MEG work reporting left ATL composition effects in a rapid 200-250 ms window (Pylkkänen, 2019a, 2019b).

One explanation for this divergence lies in the specific methodological trade-offs of our parametric design. In prioritising a parametric manipulation of specificity gradient, we did not impose the same strict controls on the lexical-statistical properties that are known to influence language and conceptual processing. Prior studies demonstrating an early composition effect carefully matched stimuli for factors like transition probability (e.g., Westerlund & Pylkkänen, 2014). In our stimulus set, transition probabilities varied considerably across items because our adjectives were selected to approximate intermediate specificity levels rather than to form highly predictable collocations. This variability may have introduced temporal jitter in compositional processing across trials, obscuring any composition-related signals that might otherwise have emerged in a consistent time window.

Additionally, several design differences may have modulated whether and when composition effects emerged. First, the global experimental contexts differed between the present study and prior work. Prior studies used binary designs that contrasted adjective-noun phrases with single words. This may have allowed participants to adopt a consistent “compositional” processing mode. Whereas, the current parametric design requires the brain to evaluate, on any given trial, one of three distinct specificity levels. This mixed, unpredictable context may discourage a rapid, automatic composition strategy but instead promote a strategy that is amenable to both words and phrases.

Second, the conceptual content of the adjectives was divergent across studies. The adjectives in previous work were often perceptible properties (such as colour, size), while the ones adopted in this study (like *nocturnal*) can be more abstract, which were chosen to approximate a specificity level between general (e.g. *bird*) and specific (e.g. *owl*) concepts. Thus, accessing and combining distinct types of concepts might engender compositional responses outside the expected 200-250 ms (Law et al., 2026).

These methodological considerations highlight boundary conditions for detecting ATL composition effects and suggest directions for future work. Systematic manipulation of transition probability, adjective type, and experimental context would help clarify which factors are necessary *versus* sufficient for eliciting rapid composition signatures. More broadly, our findings complement prior evidence for a unified semantic hub (Law et al., 2026) by revealing that *within* this unified architecture, the computational demands placed on the hub vary depending on the nature of the semantic task. Law et al. (2026) demonstrated that the bilateral ATL is sensitive to both semantic memory variables (noun concreteness) and compositional variables (adjective type), providing evidence that the same neural substrate supports both functions. The current study extends this by showing that even when the ATL operates on both word and phrasal inputs, the processing demands differ qualitatively: accessing high-specificity concepts appears to require sustained individuation processes that are not engaged to the same degree during composition.

## Data and Code Availability

The neuroimaging dataset is available at https://www.mrc-cbu.cam.ac.uk/publications/opendata/. Code and analysis pipelines are available at https://github.com/rmc-law/NocturnalBird.

## Author Contributions

R.M.C.L.: Conceptualisation, Methodology, Software, Validation, Formal Analysis, Investigation, Writing – Original Draft, Writing – Review & Editing, Visualisation, Project Administration, Data Curation

O.H.: Conceptualisation, Methodology, Resources, Writing – Review & Editing, Supervision, Project Administration, Funding Acquisition

M.A.L.R: Conceptualisation, Methodology, Resources, Supervision, Writing – Review & Editing, Project Administration, Funding Acquisition

All authors reviewed and approved the final manuscript.

## Acknowledgements

We are grateful to Clare Cook, Lucy MacGregor, Steve Eldridge, Máté Aller, Rebecca Williams, Fatih Serin, Marius Mada, and Karen Kabakulu for their assistance with data collection.

## Author notes

Open access: For the purpose of open access, the UKRI-funded authors O.H. and M.A.L.R. have applied a Creative Commons Attribution (CC BY) license to any Author Accepted Manuscript version arising from this submission.

## Declaration of the use of AI

No AI has been used in planning and carrying out the research, or writing the paper.

## Supplementary figures

**Supplementary Figure 1.**
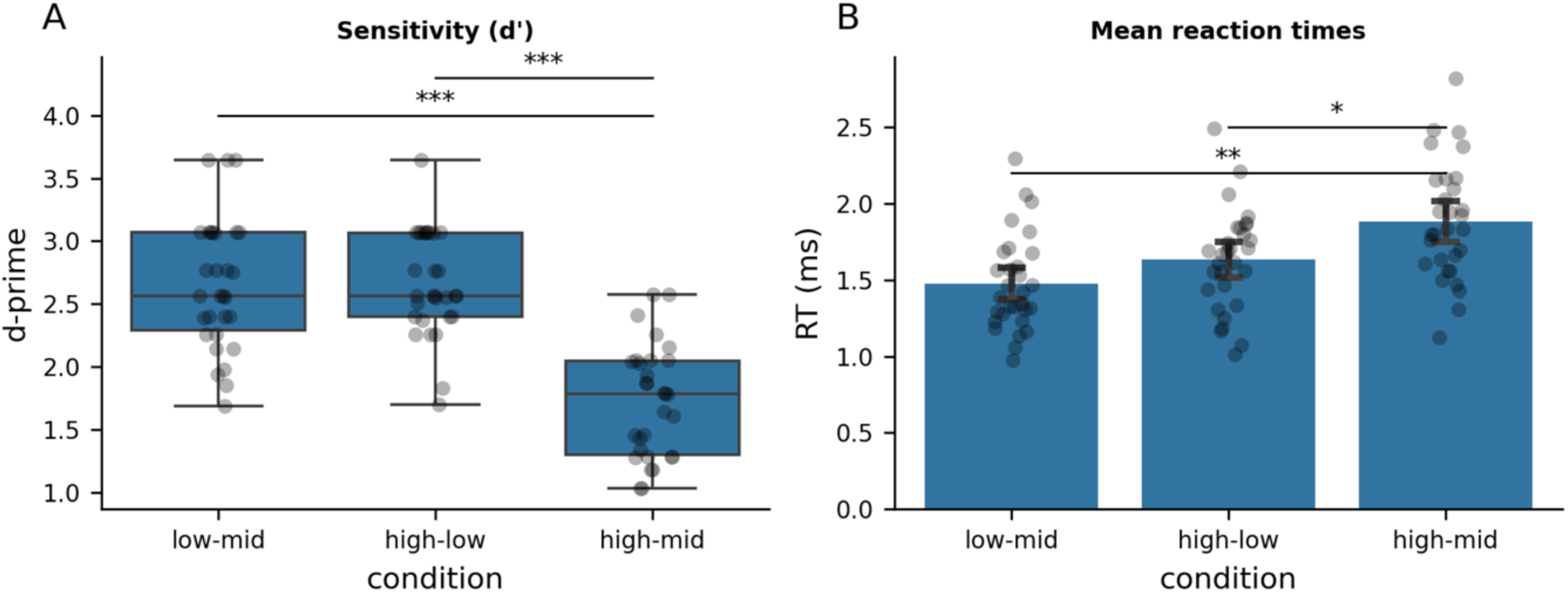
Behavioural norming results. Sensitivity (d’, A) and mean reaction time (RT, B) from the forced-choice specificity rating task across the three pairwise discrimination conditions. Boxes show the median and interquartile range; whiskers extend to 1.5× the IQR; individual points represent participant means. Asterisks denote significant pairwise differences following Tukey HSD correction (*** p < .001, ** p < .01, * p < .05).

**Supplementary Figure 2.**
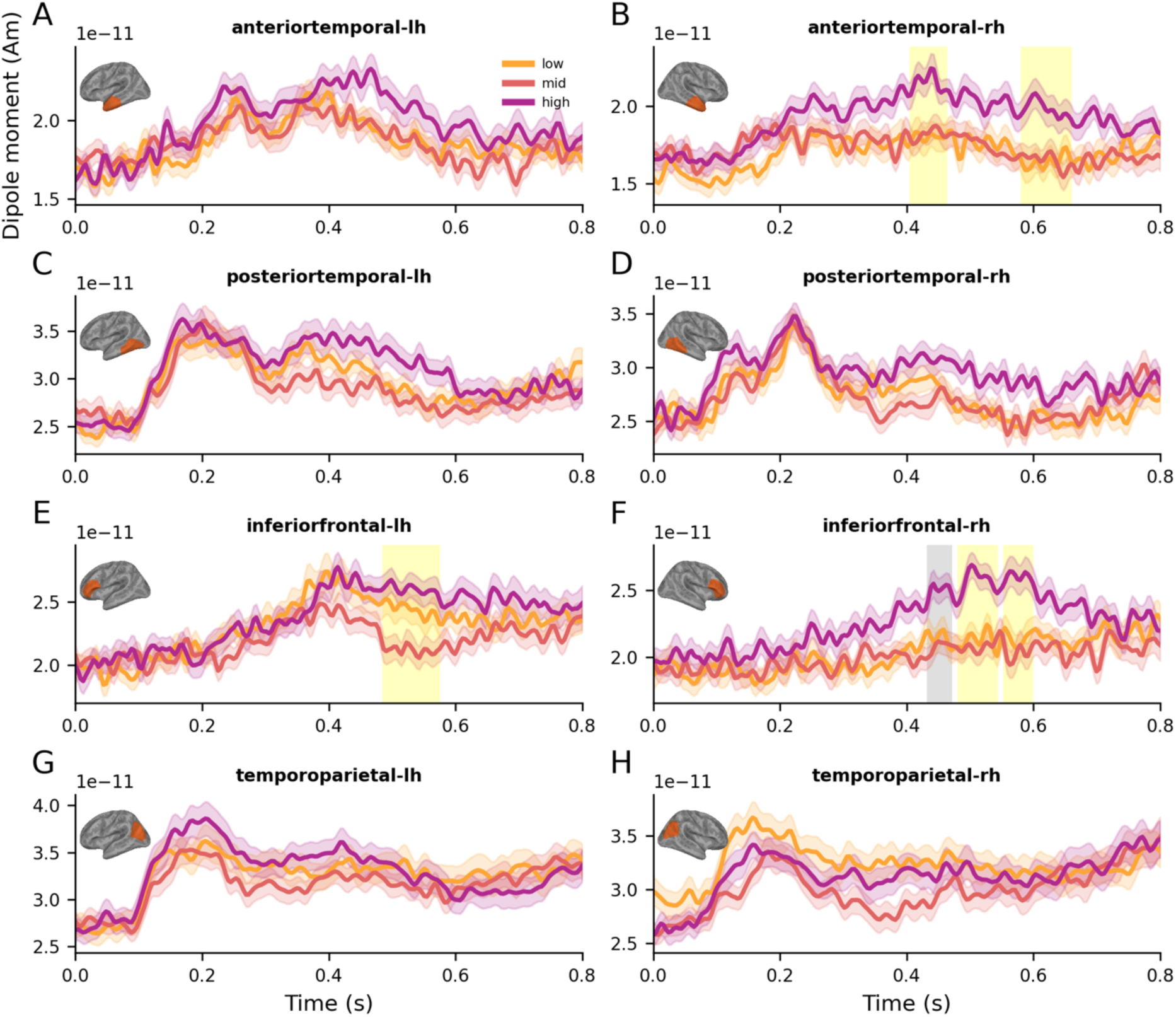
Source time courses in each tested regions of interest relative to noun onset. Time courses represent group averages of each condition and error represents one within-subjects standard error of the mean. Shaded vertical spans represent temporal clusters identified in cluster-based permutation tests: yellow indicates statistical significant effects; grey indicates marginal significance.

**Supplementary Figure 3.**
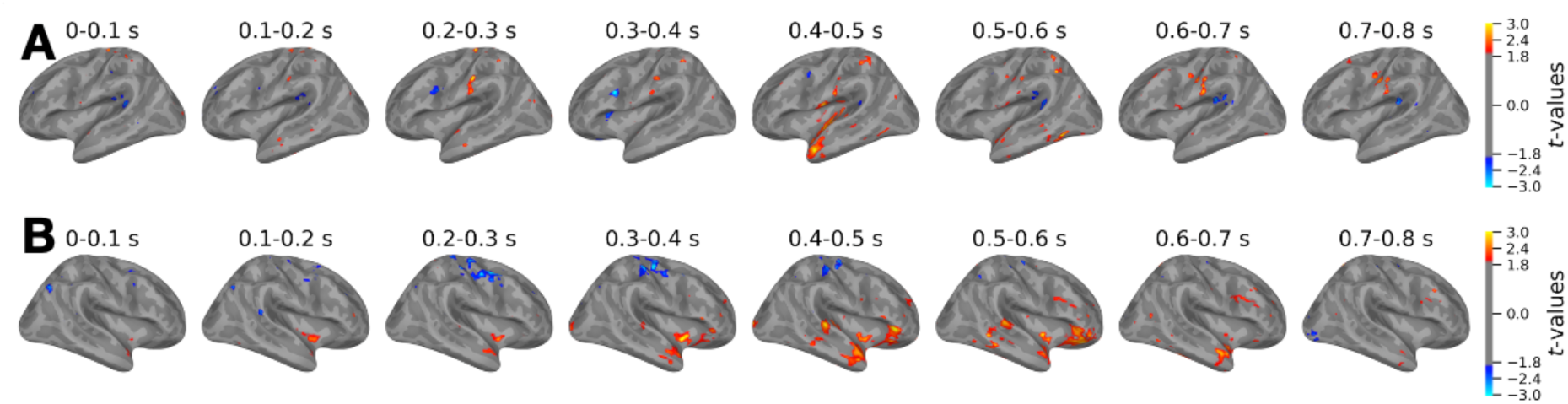
Uncorrected maps of t-values (high-low) from the onset of the noun, in 100 ms increments, in the left (A) and right (B) hemispheres.

