## Supplemental figures for "Anterior temporal lobes show route-dependent sensitivity to conceptual specificity: Evidence from MEG and EEG"

### Supplementary figures

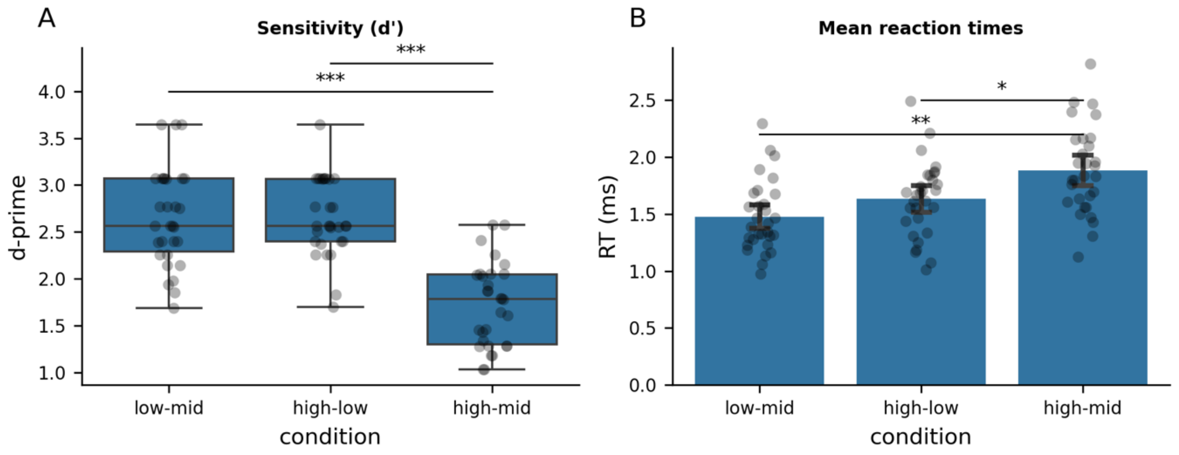


*Supplementary Figure 1. Behavioural norming results.* ***.*** *Sensitivity (d', A) and mean reaction time (RT, B) from the forced-choice specificity rating task across the three pairwise discrimination conditions. Boxes show the median and interquartile range; whiskers extend to 1.5× the IQR; individual points represent participant means. Asterisks denote significant pairwise differences following Tukey HSD correction (*** p < .001, ** p < .01, * p < .05).*

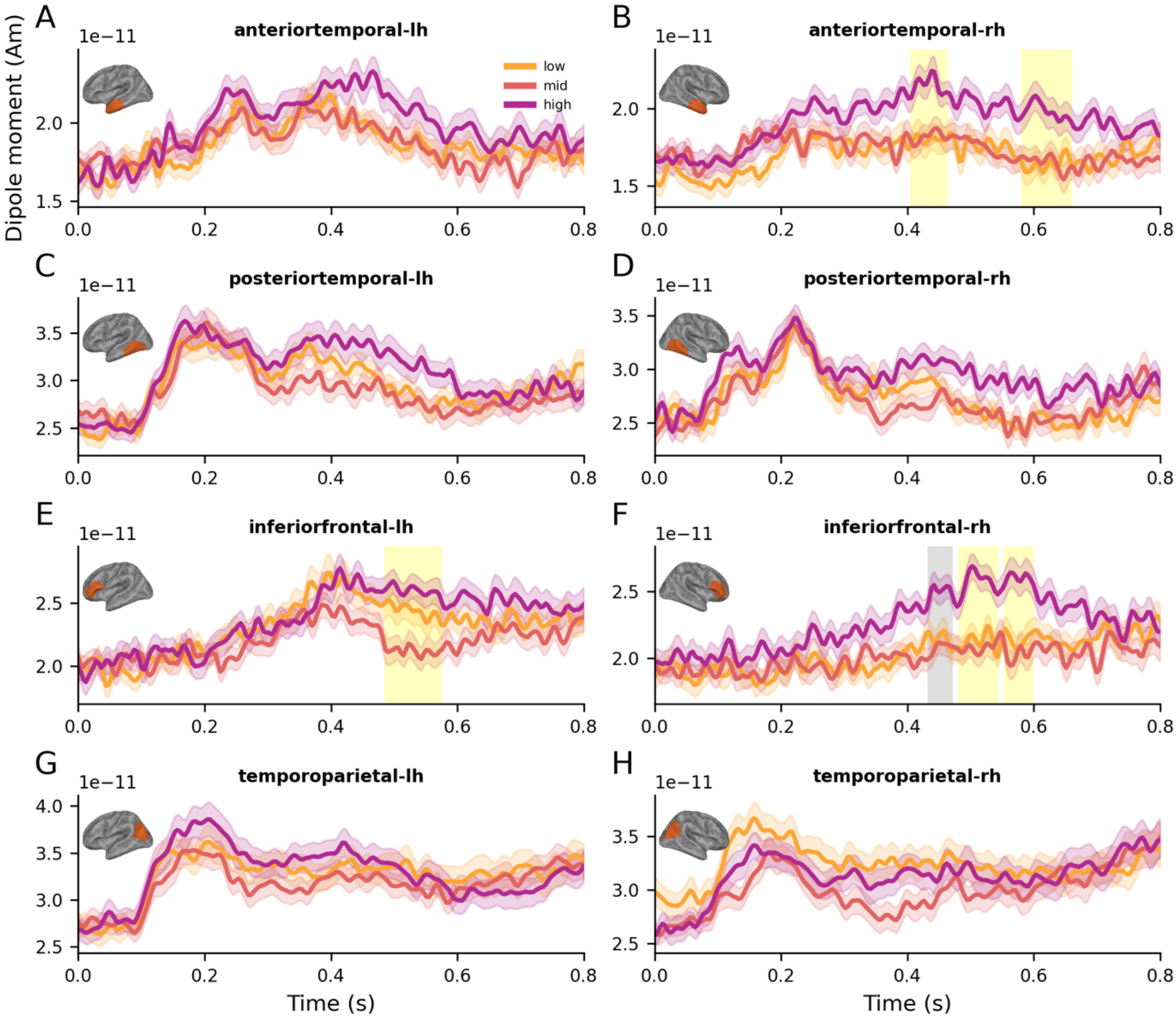


*Supplementary Figure 2. Source time courses in each tested regions of interest relative to noun onset. Time courses represent group averages of each condition and error represents one within-subjects standard error of the mean. Shaded vertical spans represent temporal clusters identified in cluster-based permutation tests: yellow indicates statistical significant effects; grey indicates marginal significance.*

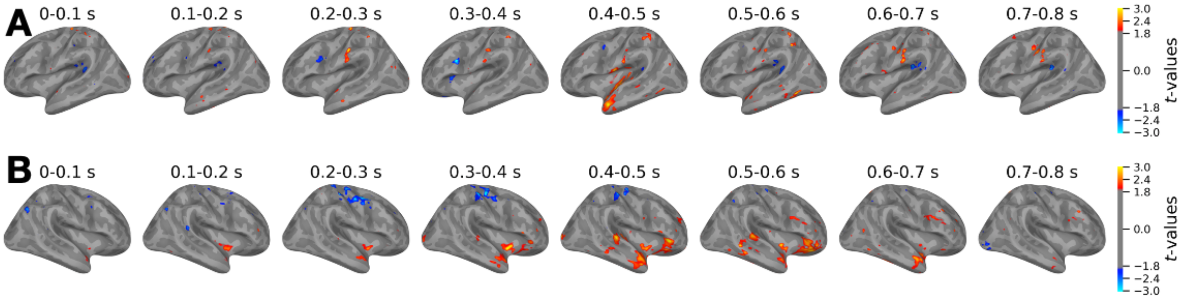


*Supplementary Figure 3. Uncorrected maps of t-values (high-low) from the onset of the noun, in 100 ms increments, in the left (A) and right (B) hemispheres.*
